# Improving the performance of supervised deep learning for regulatory genomics using phylogenetic augmentation

**DOI:** 10.1101/2023.09.15.558005

**Authors:** Andrew G Duncan, Jennifer A Mitchell, Alan M Moses

**Affiliations:** Department of Cell & Systems Biology, University of Toronto

## Abstract

**Motivation:** Supervised deep learning is used to model the complex relationship between genomic sequence and regulatory function. Understanding how these models make predictions can provide biological insight into regulatory functions. Given the complexity of the sequence to regulatory function mapping (the cis-regulatory code), it has been suggested that the genome contains insufficient sequence variation to train models with suitable complexity. Data augmentation is a widely used approach to increase the data variation available for model training, however current data augmentation methods for genomic sequence data are limited.

**Results:** Inspired by the success of comparative genomics, we show that augmenting genomic sequences with evolutionarily related sequences from other species, which we term phylogenetic augmentation, improves the performance of deep learning models trained on regulatory genomic sequences to predict high-throughput functional assay measurements. Additionally, we show that phylogenetic augmentation can rescue model performance when the training set is down-sampled and permits deep learning on a real-world small dataset, demonstrating that this approach improves experimental data efficiency. Overall, this data augmentation method represents a solution for improving model performance that is applicable to many supervised deep learning problems in genomics.

**Availability and implementation:** The open-source GitHub repository agduncan94/phylogenetic_augmentation_paper includes the code for rerunning the analyses here and recreating the figures.

## Introduction

Supervised deep learning approaches can predict chromatin accessibility^1–3^, transcription factor binding^4^, enhancer activity^5^, and other assays from genomic sequence^6^. The performance of trained deep learning models is measured using held-out test datasets, ruling out simple overfitting to limited training datasets. Aside from demonstrating powerful predictive performance, models that learn general biological features from experimental data can lead to improved understanding of biological phenomena. Indeed, the application of various post hoc techniques^7,8^ have revealed that deep learning models trained on regulatory sequences learn both known and novel motifs and motif interactions^1–5,9,10^.

Using the tools of the deep learning era^11,12^, computational biologists can now fit models with complexity that approaches the true biochemical complexity of transcriptional regulation^13,14^. However, training models with 10^6^-10^8^ parameters requires large and diverse datasets, which are available only for well-studied model organisms and human cell lines. A major challenge in the field is determining how to train more complex deep learning models for applications outside of the most data-rich systems. A proposed solution is to substantially increase data volume by performing assays on randomly generated synthetic sequences, and then evaluating models trained on these sequences using true genomic sequences^13,15^. The reasoning behind this approach is that the genome does not contain sufficient variation to learn all aspects of the cis-regulatory code. Although, this approach may not be suitable for all assays and requires additional experimentation.

Data augmentation is a technique used in deep learning to increase performance of complex models by training models on transformed versions of the input data, thereby increasing the number of training examples. This technique finds application across a variety of fields including computer vision^16^ and natural language processing^17^. However, there are limited domain-specific augmentation techniques for genomic sequence data. Two commonly used augmentation approaches include reverse complements^18^ and genomic shifts^5,19^. Recently, sequence augmentations inspired by evolution, including point mutations, inversions, and deletions, have shown promise in improving supervised model performance and interpretability^20^. However, because these approaches introduce random sequence modifications to simulate evolution, they do not account for any functional constraint on the sequence.

Homologous sequences (or homologs for short) are genomic sequences from different species that share a common ancestor and may be under functional constraint, but have likely diverged in primary sequence^21,22^. Previously, homologs have been proposed as a form of sequence augmentation for self-supervised learning, as they can be thought of as transformed versions of their ancestral sequence that retain the same biological function (analogous to augmentations that do not alter the semantic meaning in images)^23–25^. Additionally, it has been demonstrated that training deep learning models on functional genomic assays from both human and mouse can improve model performance compared to using data from a single species^25^, indicating there is a benefit to training models on sequences from multiple species. Here, we investigate the use of homologs as a data augmentation method for supervised deep learning models that predict functional genomic assays from genomic sequence. We find that augmenting genomic sequences with homologs can improve the performance of supervised deep learning models. Furthermore, we show that phylogenetic augmentation can restore performance on models trained on down-sampled training sets, indicating that this approach has the potential to enhance data efficiency when training size is low. We then apply phylogenetic augmentation to a real-world small genomic dataset, where it allows for the training of a deep learning model.

## Results

### Phylogenetic augmentation: A method for augmenting genomic sequences using multi-species genome alignments

We define phylogenetic augmentation for supervised deep learning as the transformation of a genomic sequence from one species into a homolog from another species. This approach leverages homologs extracted from multi-species genome alignments to improve the diversity of input training data. By presenting these homologs as augmented versions of training sequences, deep learning models see a broader array of sequences during the training process.

The application of phylogenetic augmentation within supervised deep learning problems involves three phases, as illustrated in Figure 1A. Prior to model training, an initial preprocessing step is performed to extract homologs for each genomic sequence in the training set from a multi-species genome alignment that contains the species being investigated (Figure 1B). This is done before training, so that existing alignment tools can be used to extract homologs (see methods). During model training, phylogenetic augmentation is applied to all training sequences at batch generation (Figure 1C). This significantly increases the number of training examples that the model encounters. Following a previous approach, after training, the models are fine-tuned on the original genomic sequences to further improve performance to reduce potential bias from including functionally diverged homologs^20^.

**Figure 1.**
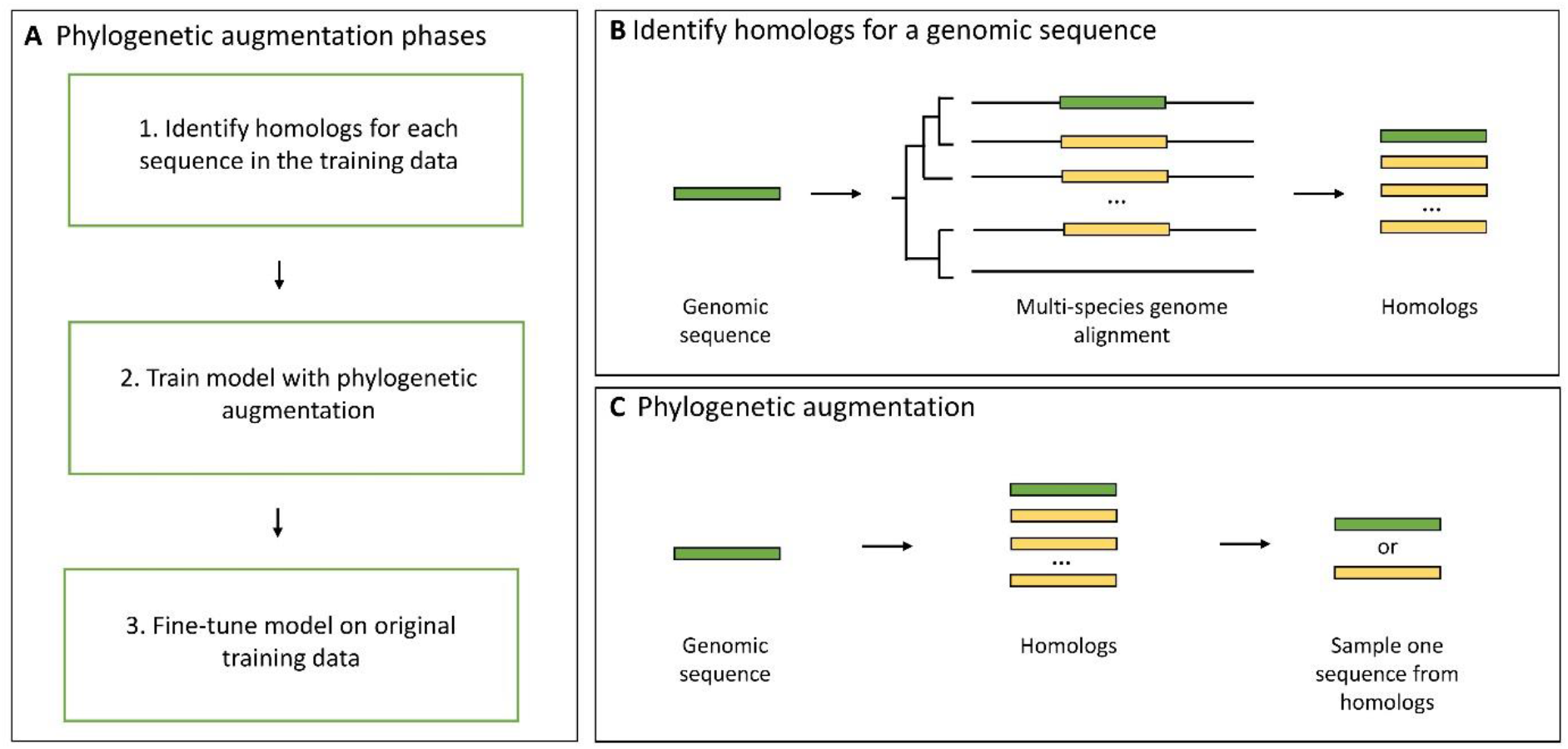
An overview of the phylogenetic augmentation method. (A) The three phases of phylogenetic augmentation for model training. (B) During preprocessing, homologs (yellow boxes) of each training sequence (green boxes) are identified in different genomes (black lines) using multi-species genome alignments. (C) Phylogenetic augmentation is implemented as the transformation of a genomic sequence to a random homologous sequence.

### Phylogenetic augmentation improves CNN prediction performance on held-out test sets

To investigate whether phylogenetic augmentation used on a supervised deep learning problem improved model performance, we trained convolutional neural networks (CNNs) with phylogenetic augmentation to predict *Drosophila* S2 STARR-seq activity driven by a housekeeping or developmental promoter^5^, and compared performance to a baseline where no phylogenetic augmentation or fine-tuning was applied. For each sequence in the training set, homologs for 136 species in the *Drosophila* genus were extracted from a multi-species genome alignment. To explore the effect of CNN architecture, three CNN architectures of varying complexities were trained on this data (DeepSTARR^5^, ExplaiNN^9^, and Motif DeepSTARR), and their performance was measured on a held-out test set (see methods).

We observed that all CNN models demonstrated an increase in test set performance compared to the baseline with the inclusion of phylogenetic augmentation and fine-tuning (Figure 2A). For example, the average DeepSTARR model performance (Pearson correlation coefficient; PCC) increased from 0.661 (± < 0.01 S.E.) to 0.689 (± < 0.01 S.E.) (+4.2%) for the developmental enhancer activity, and 0.741 (± < 0.01 S.E.) to 0.779 (± < 0.01 S.E.) (+5.1%) on the housekeeping enhancer activity (grey and blue points). The ExplaiNN^9^ CNN had a smaller performance increase than the other two architectures. This can be attributed to differences between the ExplaiNN and the DeepSTARR models. ExplaiNN uses linear combinations of learned motif representations to make predictions, and unlike the other two CNN models which have fully connected layers, cannot learn complex non-linear interactions between different transcription factor motifs^9^. While fine-tuning the models on the original training data after phylogenetic augmentation further improved performance (green points and blue points), fine-tuning the baseline models was not sufficient by itself to achieve the same level of performance increase seen with phylogenetic augmentation and fine-tuning (yellow points and blue points).

**Figure 2.**
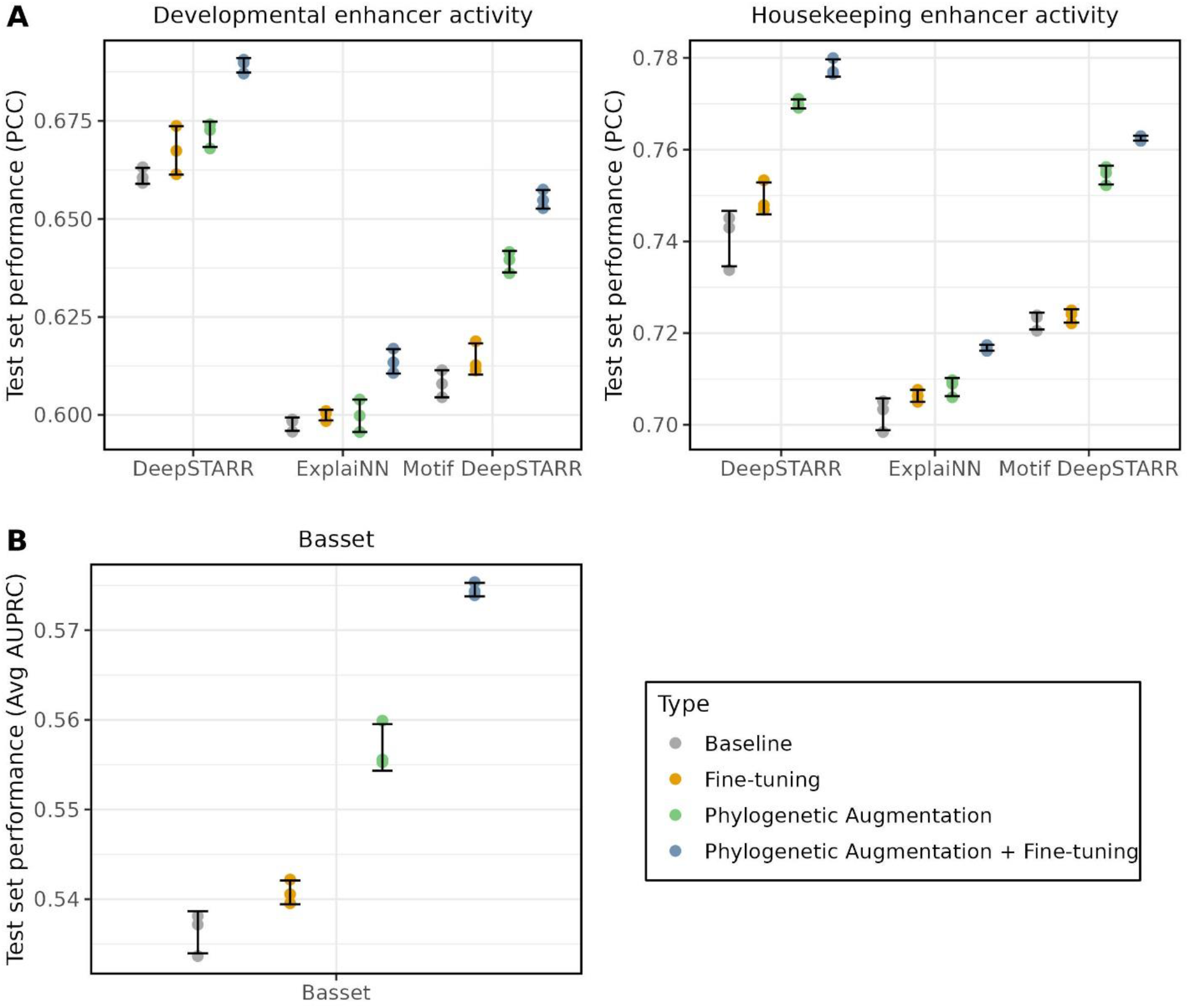
Phylogenetic augmentation improves model performance with various CNN architectures. (A) CNN test performance (Pearson correlation coefficient; PCC) is shown on the y-axis for the *Drosophila* S2 Developmental (left) and Housekeeping (right) enhancer activity for trained models. (B) Basset model test performance (average area under the precision recall curve; avg AUPRC) is shown on the y-axis for 164 cell-type specific chromatin accessibility (DNase-seq) experiments for trained models. (A-B) The grey points are the baseline models, which were trained with no phylogenetic augmentation or fine-tuning. The yellow points are the baseline models that have been fine-tuned on the original data. The green points are models trained with phylogenetic augmentation. The blue points are the models trained with phylogenetic augmentation that have been fine-tuned on the original data. The black error bars represent the standard deviation of the three replicates for each CNN architecture and dataset.

Next, we performed a similar analysis using the Basset model to predict binary DNase-seq peaks across 164 human cell-lines, which we refer to as the Basset dataset^1^. For every sequence in the training set, homologs from a clade of 43 species that included *Homo sapiens* were extracted from a mammalian genome alignment. Again, we observed improved model performance on a held-out test set when comparing the baseline models to the models with phylogenetic augmentation and fine-tuning, with the average area under the precision recall curve (AUPRC) increasing from 0.536 (± < 0.01 S.E.) to 0.575 (± < 0.01 S.E.) (+7.2%) (Figure 2B). Together, these results demonstrate that phylogenetic augmentation is a useful data augmentation approach for training supervised deep learning models on genomic sequences, though the magnitude of improvement is dependent on model architecture.

### Phylogenetic augmentation improves data efficiency in supervised deep learning

Some regulatory datasets may not include enough regions of interest for effective machine learning, which could lead to overfitting on training data. Phylogenetic augmentation could serve as a method for improving data efficiency when training models on smaller genomic datasets. To investigate this, we down-sampled different fractions of the training sequences for the *Drosophila* S2 STARR-seq dataset and applied phylogenetic augmentation during DeepSTARR model training to determine if test set performance could be rescued. For each fraction of the training sequences, phylogenetic augmentation with fine-tuning improved test set performance compared to the baseline (blue and grey points) (Figure 3A). At 40% and 20% of the original training sequences, phylogenetic augmentation plus fine-tuning was sufficient to rescue the baseline model’s performance on the test set for the developmental and housekeeping enhancer activities, respectively (Figure 3A, dotted grey lines). The largest performance improvements were seen when phylogenetic augmentation and fine-tuning was applied to only 10% of the dataset, with gain over the baseline model of 0.133 for the developmental enhancer activity compared to an improvement of 0.0318 on 100% of the original dataset (Figure 3A). A similar experiment was performed for the Basset dataset using the Basset model, with phylogenetic augmentation plus fine-tuning again improving test-set performance for all fractions tested (Figure 3B). At around 40% of the original training data, phylogenetic augmentation was able to rescue the baseline model performance seen with all the training data (Figure 3B, dotted grey line). These results indicate that phylogenetic augmentation can enhance the data efficiency of supervised deep learning models, enabling them to make better predictions with less data.

**Figure 3.**
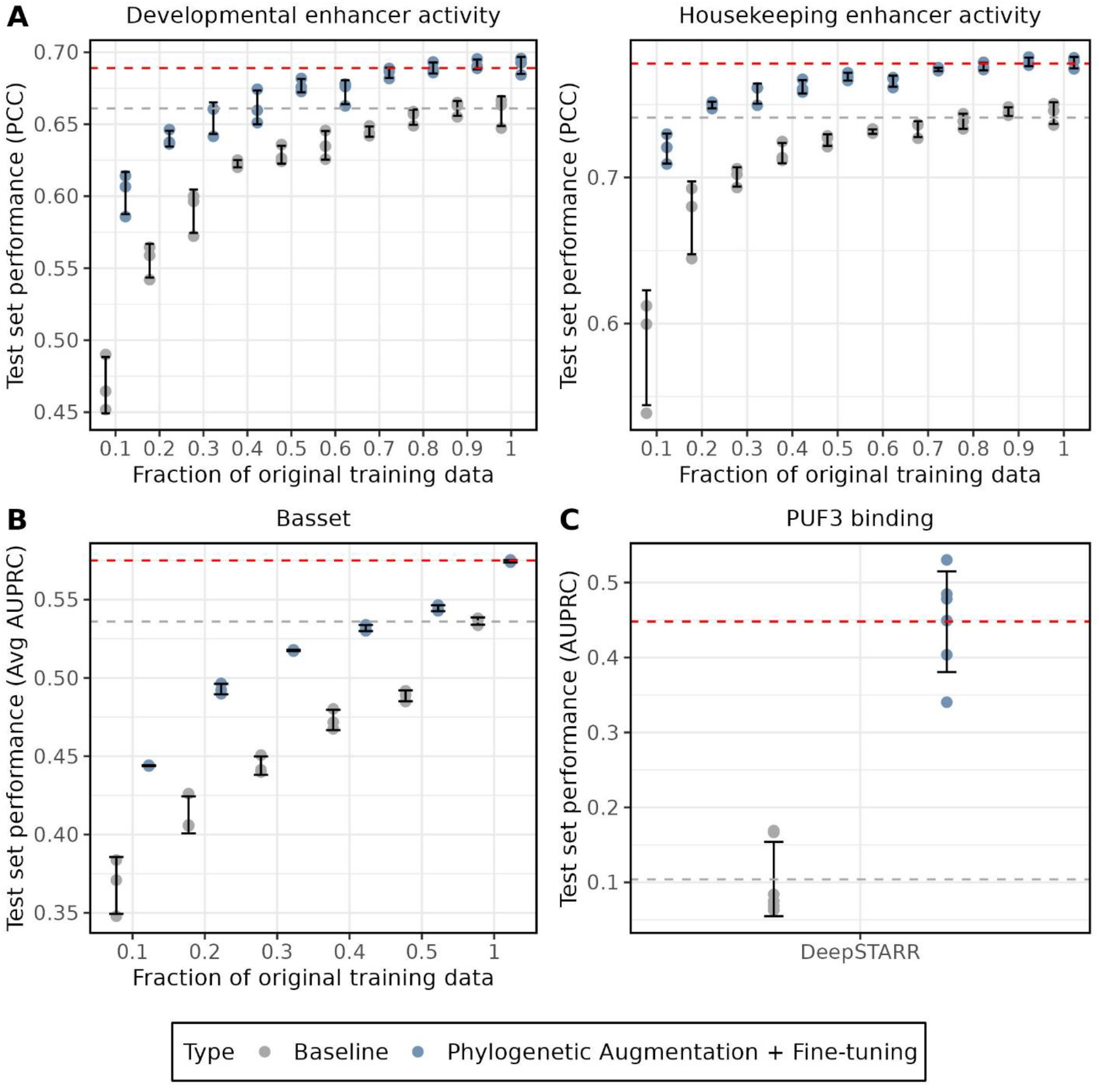
Phylogenetic augmentation improves data efficiency when training size is reduced. (A) DeepSTARR test performance (Pearson correlation coefficient; PCC) is shown on the y-axis for the *Drosophila* S2 Developmental (left) and Housekeeping (right) enhancer activity for trained models. (B) Basset test performance (average area under the precision recall curve; avg AUPRC) is shown on the y-axis for 164 cell-type specific chromatin accessibility (DNase-seq) experiments for trained models. (A-B) The x-axis represents the fraction of the original training data that was sampled during model training. (C) The DeepSTARR test performance (area under the precision recall curve; AUPRC) is shown on the y-axis for the binary classification prediction of *S. cerevisiae* 3’UTR PUF3 binding. (A-C) The grey points are the baseline models, which were trained with no phylogenetic augmentation or fine-tuning. The blue points are the models trained with phylogenetic augmentation that have been fine-tuned on the original training data. The black error bars represent the standard deviation of the replicates for each CNN architecture and dataset. The dotted grey line represents the average test performance on the original training data with no phylogenetic augmentation or fine-tuning. The dotted red line represents the average test performance on the original training data with phylogenetic augmentation and fine-tuning.

Due to the large performance increases seen on the down-sampled *Drosophila* dataset, we next asked whether phylogenetic augmentation could be applied to a real-world example of a small dataset where it is challenging to train a complex deep learning model. We chose to predict the binding of the RNA-binding protein PUF3 to 3’ untranslated regions (3’UTRs) of *Saccharomyces cerevisiae*^26^ because this dataset is more than an order of magnitude smaller than the *Drosophila* dataset (∼5,000 *S. cerevisiae* 3’UTRs vs ∼200,000 *D. melanogaster* STARR-seq regions). The hyperparameters controlling the complexity of the DeepSTARR model were optimized for the *D. melanogaster* S2 STARR-seq dataset, and the model contains ∼400,000 parameters. We reasoned that a model of this complexity would be challenging to train on the *S. cerevisiae* 3’UTRs.

To test this, we trained a DeepSTARR model as above, and as expected, found that the DeepSTARR model had a test classification performance (AUPRC) of 0.104 (± < 0.1 S.E.) on the test data (Figure 3C), only marginally better than the test performance of 0.0427 (± < 0.01 S.E.) seen on a scrambled label control. Remarkably, when phylogenetic augmentation and fine-tuning was applied to this dataset during model training, the average test performance increased by over 4-fold to 0.448 (± < 0. 1 S.E.) (Figure 3C). To test the biological relevance of this performance increase, we wondered whether the large test performance increase could be explained by the augmented model learning the known PUF3 consensus motif^27^. We performed global importance analysis^8^ of the PUF3 motif for both the baseline model and the model with phylogenetic augmentation (see methods). Only with phylogenetic augmentation does the model place importance on the PUF3 motif (Supplementary figure 1). Taken together, these results demonstrate that phylogenetic augmentation can enable training of complex deep learning models and learning of biologically relevant features on small genomic datasets.

### Exploring hyperparameters for phylogenetic augmentation

To assess the impact of different hyperparameters on phylogenetic augmentation, we trained multiple DeepSTARR models on the *Drosophila* S2 STARR-seq dataset and varied the number of species used and the rate at which phylogenetic augmentation was applied.

First, we examined how the number of species and their total evolutionary distance used during phylogenetic augmentation impacted model test performance. We arranged the *Drosophila* species from the *Drosophila* multi-species alignment based on increasing evolutionary distance from *D. melanogaster*. Then, we trained multiple DeepSTARR models with phylogenetic augmentation and fine-tuning, progressively incorporating more distant species from which to draw homologs. We observed that including homologs for just one additional species yielded a noticeable performance increase for both enhancer activity measurements (Figure 4A). Interestingly, model test performance improvements plateaued around 10 species for housekeeping enhancer activity, but for developmental enhancer activity performance started decreasing after 10 species. Performing the same analysis starting with the most distant species had minimal test set performance increases for the first 20 species, though there was some improvement over the baseline model (Supplementary figure 2). These results suggest that a handful of closely related species is sufficient for improving model test performance through phylogenetic augmentation, and that including too many distant species may decrease phylogenetic augmentation performance gains in some cases.

**Figure 4.**
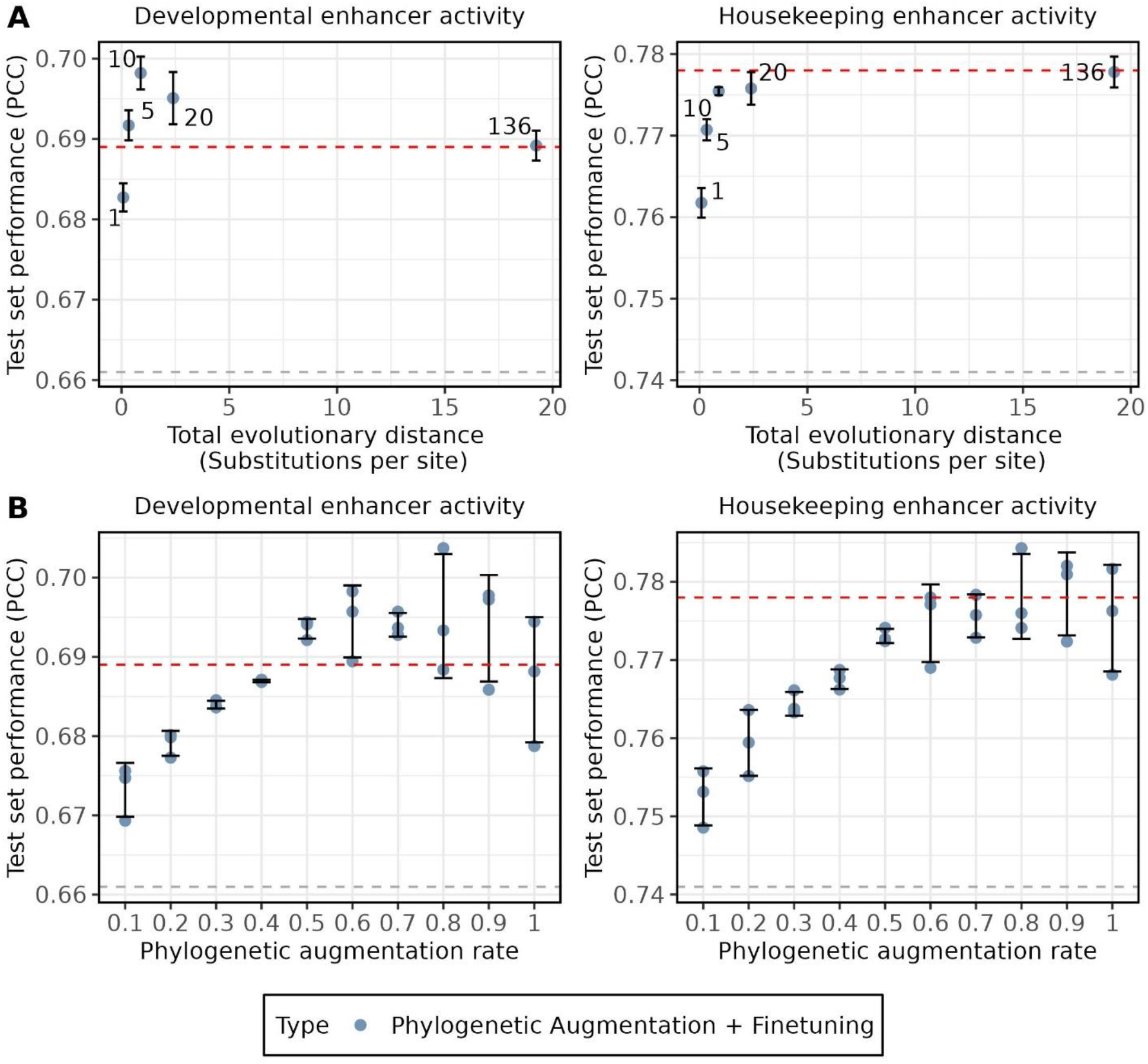
Hyperparameters for phylogenetic augmentation impact test set performance. (A) DeepSTARR test performance (Pearson correlation coefficient; PCC) is shown on the y-axis for Developmental (left) and Housekeeping (right) enhancer activity for trained models. The x-axis represents the total evolutionary distance of species included during phylogenetic augmentation, measured using substitutions per site. The labels represent the total number of species used to train each model. The blue dots represent the average test set performance across replicates. (B) DeepSTARR test performance (Pearson correlation coefficient; PCC) is shown on the y-axis for Developmental (left) and Housekeeping (right) enhancer activity for trained models. The x-axis represents the rate at which phylogenetic augmentation is applied during model training. The blue dots represent the test set performance for individual replicates. (A-B) The black error bars represent the standard deviation of the three replicates that were trained for each number of species. The dotted grey line represents the average test performance on the original training data with no phylogenetic augmentation or fine-tuning. The dotted red line represents the average test performance on the original training data with phylogenetic augmentation and fine-tuning using all 136 *Drosophila* species and a phylogenetic augmentation rate of 1.

Next, we explored how the rate at which phylogenetic augmentation is applied during training affects test performance. We trained DeepSTARR models on the *Drosophila* S2 STARR-seq dataset, applying different rates of phylogenetic augmentation at batch generation. For example, a rate of 0.5 signified that 50% of the sequences in each batch underwent phylogenetic augmentation. This was carried out using homologs for all 136 *Drosophila* species from the *Drosophila* multi-species alignment. As shown in Figure 4B, an increase in the rate of phylogenetic augmentation led to improved test set performance for both enhancer activity measurements. However, past a rate of 0.5, test set performance became more variable and subsequently decreased or plateaued. We hypothesized that the performance variation was caused by the higher phylogenetic augmentation rates adding too many sequences with diverged function. Consistent with this hypothesis, the effect was reduced when we repeated the analysis with the 10 closest species (Supplementary figure 3). These results indicate the importance of the rate of phylogenetic augmentation when using this augmentation approach, and that it is not always best to apply this augmentation to every sequence in a batch.

## Discussion

Here, we introduced a data augmentation method for supervised deep learning of genomic sequences that takes advantage of multi-species genome alignments and the phylogenetic relationship between homologous sequences. This approach improved the performance of regression and classification problems for three different functional genomic datasets across two different kingdoms of life, indicating it is a general approach for augmenting genomic sequence data. Additionally, we used phylogenetic augmentation on a reduced dataset to rescue model performance seen with the original training dataset, demonstrating that this approach improves the data efficiency of models.

In machine learning, data augmentation finds common use when datasets are too small such that deep learning models will memorize or overfit the training data. Although many regulatory datasets, such as those resulting from functional genomic experiments, contain large quantities of data, not all do. For instance, curated databases like the VISTA enhancer database include only a few thousand mouse and human enhancers^28^. Potentially, data augmentation approaches like phylogenetic augmentation could contribute to addressing the challenge of training deep learning models on these small genomic datasets. Our results suggest that phylogenetic augmentation is most effective on small datasets (Figure 3A). In line with this observation, applying phylogenetic augmentation to a DeepSTARR model trained on PUF3 binding across *S. cerevisiae* 3’UTRs resulted in a substantial increase in test set performance over the baseline model (Figure 3C) and the ability to learn the PUF3 consensus motif (Supplementary figure 1).

Regulatory elements, like promoters and enhancers, undergo evolution at different rates, which can lead to varying rates of element turnover^29^. Thus, the efficacy of this approach can be influenced by the species chosen for augmentation, as some elements may lose function quicker than others. Since the *Drosophila* S2 STARR-seq data contains enhancers, which have higher rates of turnover, we restricted species to those in the *Drosophila* genus. Had we included only species outside of the genus, such as other insects, the number of homologs identified would have been noticeably lower. Consequently, the phylogenetic augmentation method likely would have had a decrease in effectiveness. We observed that even for the same type of regulatory elements, there can be differences in the effectiveness of distant species. Test performance increases from applying phylogenetic augmentation to the *Drosophila* S2 developmental enhancers peaked at 10 species before slowly falling (Figure 4A). Meanwhile, test performance of the housekeeping enhancers continued to increase (albeit minimally) with the number of species (Figure 4A). A potential explanation is that the expression of housekeeping genes is more conserved than tissue specific genes^30^, suggesting housekeeping enhancers may be under stronger functional conservation. Based on these results, we recommend using 10 closely related species as a starting point when applying phylogenetic augmentation during model training.

While it has been demonstrated that training a CNN to predict both mouse and human functional genomic experiments from DNA sequence can improve predictive performance on an individual species^25^, this approach requires data from multiple assays which require additional time, resources, and expertise. While there may be data from relevant experimental assays available, these are often restricted to a handful of well-studied species (e.g., ENCODE Project). Our approach does not have these limitations, as it only requires sequenced genomes and whole genome alignments. In the case where there are no available multi species genome alignments that include a species of interest, evolutionary augmentation using simulated mutations can be applied^20^, or alignments can be generated from resources such as the Earth Biogenome Project^31^ or the NCBI genome database^32^. We recommend creating custom Cactus alignments^33^ using HAL^34^, which was designed to work with large multi species genome alignments.

As researchers are increasingly relying on the features learned by deep learning models trained on sequence data to obtain biological insights of regulatory function, it is crucial that these models are not overfitting to their training data and generalize to unseen sequences. The method we proposed here holds the potential for application in any supervised deep learning problem that uses non-coding genomic sequences as input, even when the number of training examples is insufficient for complex deep learning models. With the increasing availability of large alignments such as the 240-way mammalian alignment from Zoonomia^22^, the 239-way primate alignment from Zoonomia^35^, the 341-way avian alignment from B10K^36^, and the 168-way *Drosophila* alignment used here (Supplementary table 5), our approach is widely applicable across a variety of species.

## Methods

### Datasets

We used published data from a massively parallel reporter assay (MPRA) experiment measuring enhancer activity across the *Drosophila Melanogaster* genome in the S2 cell-line^5^. The MPRA was performed once using a housekeeping promoter and once with a developmental promoter to investigate enhancer-promoter dependencies, resulting in two measurements per sequence. The dataset consists of 7,062 and 11,658 enhancers identified in the housekeeping and developmental experiments, respectively. Additionally, 223,306 regions with varying activity levels were also included to provide sequences with a range of activity levels. All sequences are of length 249bp. To match the original manuscript, chromosome 1 was split in half, with the first half used for validation and the second half used for testing. The remainder of the data was used for model training. Instead of including reverse complements in the dataset, reverse complements were randomly applied during model training to 50% of sequences.

The Basset dataset contains 2,071,886 DNase-seq peaks from 164 different human cell-lines, along with a binary class for whether each peak is open in each cell-type. After filtering out sequences with Ns (unknown nucleotides), this was reduced to 2,021,532 regions. All sequences are of length 600bp. To match the original manuscript, the data was split into training, validation, and testing using 93%, 3.5%, and 3.5% of the data, respectively.

The yeast 3’UTR dataset contains binary RNA-binding data for the PUF3 RNA-binding protein to 4,293 3’UTRs from *S. cerevisiae*^37^ that we obtained for a previous analysis^26^. We selected PUF3 because in a previous analysis we found that PUF3 had the strongest motif signal of the 74 RNA-binding proteins from AtTract^38^ in *S. cerevisiae* 3’UTRs^26^. These 3’UTR sequences were defined as regions 200bp downstream of the stop codon of a gene. Due to the limited number of total positives (198) in the dataset, we did not use a validation set to optimize parameters. Chromosome 4 (ChrYD) was used for the testing dataset, and the remaining chromosomes were used for model training.

### Models

For the *Drosophila* S2 analysis, we used three different CNN architectures designed for DNA sequences and implemented them in TensorFlow: DeepSTARR^5^, ExplaiNN^9^, and a custom architecture we call Motif DeepSTARR. These architectures are meant to represent various design choices used for deep learning of genomic sequences. The input to these models is 249bp DNA sequences that are one-hot encoded, and the task is a regression on the MPRA enhancer activity for each promoter MPRA experiment. A summary of the encoder component of the architectures is provided below.

- **DeepSTARR**: A CNN composed of four convolutional layers followed by two fully connected layers. The convolutional layers are meant to capture sequence motifs, and the fully connected layers are meant to capture non-linear relationships between motifs.
- **Motif DeepSTARR**: A single convolutional layer with 256 filters, followed by two fully connected layers. The convolutional layer has a large filter size (19) in order to learn full motifs. The fully connected layers capture non-linear relationships between motifs.
- **ExplaiNN**: A linear combination of 256 motif representations. The motif representations are defined as a convolutional layer with 1 filter, followed by two fully connected layers.

For the Basset analysis, we used only the Basset model^1^. The input to this model is 600bp DNA sequences that is one-hot encoded, and the task is 164 multi-label binary classification on 164 different human cell-lines. A summary of the encoder component of the architectures is provided below.

- **Basset**: A CNN composed of three convolutional layers followed by two fully connected layers. The inspiration for the DeepSTARR architecture.

For the yeast 3’UTR analysis, we used the same DeepSTARR encoder that was used with the *Drosophila* S2 analysis, however the input to the model was changed to 200bp one-hot encoded DNA sequences. The task is binary classification of PUF3 binding, with 1 representing binding and 0 representing no binding.

### Identifying homologous sequences from a multi-species genome alignment

For the *Drosophila* S2 analysis, we obtained an unpublished multi-species genome alignment of 168 genomes containing 137 species from the *Drosophila* genus, including *D. melanogaster* (Supplemental data file 1). Homologs from other *Drosophila* species were extracted from the alignment for each *D. Melanogaster* training sequence using HAL^34^ and the HALPER tool^39^. If multiple homologs were found in a target organism, the first match was arbitrarily selected. The selected sequences were then resized to 249bp from the midpoint to match the length of the training sequence. This resulted in 10,582,901 homologs in total across all 191,109 sequences in the training set.

For the Basset analysis, we used the Zoonomia multi-species genome alignment of 241 mammalian species^22^. Homologs from 42 species were used for this analysis and were restricted to clades around the *Homo sapiens* node (Supplemental data file 2). The selected sequences were then resized to 600bp from the midpoint to match the length of the training sequence. This resulted in 62,538,898 homologs in total across all 1,880,029 sequences in the training set.

For the yeast 3’UTR analysis, we used a previously collected dataset of homologs from Ensembl^40^ for 24 different yeast species (Supplemental data file 3)^26^. Unlike the multi-species genome alignment approach that we had used to identify homologs for the *D. melanogaster* and *H. sapiens* sequences in the previous analyses, we used a genic approach to identify the *S. cerevisiae* homologs. Briefly, the Ensembl REST API^41^ was used to retrieve one-to-one orthologs for each gene along with the 200bp downstream of the stop codon of the corresponding gene. This resulted in a total of 71,498 homologs across all 3,721 sequences in the training set.

### Model training and fine tuning

For the *Drosophila* analysis, models were trained using TensorFlow^11^ with the Adam optimizer^42^ for 100 epochs with a learning rate of 2×10^-3^, mean squared error loss, and early stopping of 10. During model training, phylogenetic augmentation was applied to each training sequence (phylogenetic augmentation rate = 1.0 unless otherwise specified). This was done by sampling from the set of homologs for a given training sequence. A reverse complement augmentation was also applied randomly to sequences, including when no phylogenetic augmentation was used. As in a recent study of sequence augmentations, after training the models using the phylogenetic augmentation approach, we fine-tuned for 5 epochs at a lower learning rate (1×10^-4^) using only the original training sequences^20^. Phylogenetic augmentation is never applied to the validation or testing sets. For the sampling experiment, we trained the DeepSTARR model with sampled fractions of the training data. Sampling was performed separately for each replicate. We use Pearson Correlation Coefficient (PCC) to measure each model’s performance for each task on the same held-out test dataset.

The Basset models were trained similarly to the *Drosophila* analysis, with the following changes. Models were trained with the Adam optimizer for 20 epochs with a learning rate of 2×10^-3^ and binary cross entropy loss. The average area under the precision recall curve (AUPRC) was used to measure the model’s performance for each task on the held-out test dataset. This was used in place of the average area under the curve (AUC), as the data has a large class imbalance between positives and negatives.

The yeast models were trained similarly to the *Drosophila* analysis, with the following changes. Models were trained with the Adam optimizer for 50 epochs. No early stopping on the validation set was done, as there was no validation set. The area under the precision recall curve (AUPRC) was used to measure the model’s performance for the classification task on the held-out test dataset. Due to the small size of the dataset, 6 replicates were trained for each model. Additionally, as RNA is single stranded, reverse complements were not applied. As a negative control, the model was trained on the training data with the labels scrambled.

#### Global importance analysis of PUF3 motif

A set of 1,000 sequences of length 200bp were randomly generated to use as a background set for the analysis. The PUF3 consensus motif (TGTAAATA)^27^ was randomly inserted once into each background sequence to create a set of sequences with the PUF3 motif. As a control, we also created a set of sequences with a scrambled PUF3 consensus motif randomly inserted. A replicate of the baseline model and a replicate of the phylogenetic augmentation with fine-tuning model were used to predict the class of each sequence.

### Investigating hyperparameters

For the number of species analysis, the *Drosophila* phylogenetic tree was extracted from the *Drosophila* multi-species Cactus alignment^43^ using the halStats command (--tree) from the HAL package^34^. The ETE Toolkit^44^ get_distance function was used to determine the distance between *D. melanogaster* and all other species in the phylogenetic tree. These species were then sorted by their ascending evolutionary distance. The DeepSTARR model was then trained using phylogenetic augmentation on the homologs for the closest species, then the two closest species, and so on. For each set of species, three DeepSTARR replicates were trained. To measure the total evolutionary distance from *D. Melanogaster* for each set of species, the ETE Toolkit prune function (preserve_branch_length=True) was used to prune all but the current set of species from the species tree. Total distance was then calculated by summing all the branch lengths in the pruned Newick tree. The branch lengths are measured using substitutions per site of the 4-fold degenerate sites of BUSCO genes^45^.

For the phylogenetic augmentation rate analysis, the DeepSTARR model was trained on the *Drosophila* S2 dataset using increasing rates of phylogenetic augmentation. Homologs for all 136 *Drosophila* species from the alignment were used. Three replicates were run for each rate of augmentation.

Training of Basset models took substantially more time due to the length of the sequences and the dataset being an order of magnitude larger than the DeepSTARR data. Therefore, hyperparameters were not explored in Basset. Additionally, we did not investigate hyperparameters for the yeast data due to the limited data size.

#### Supplementary figures

**Supplementary figure 1.**
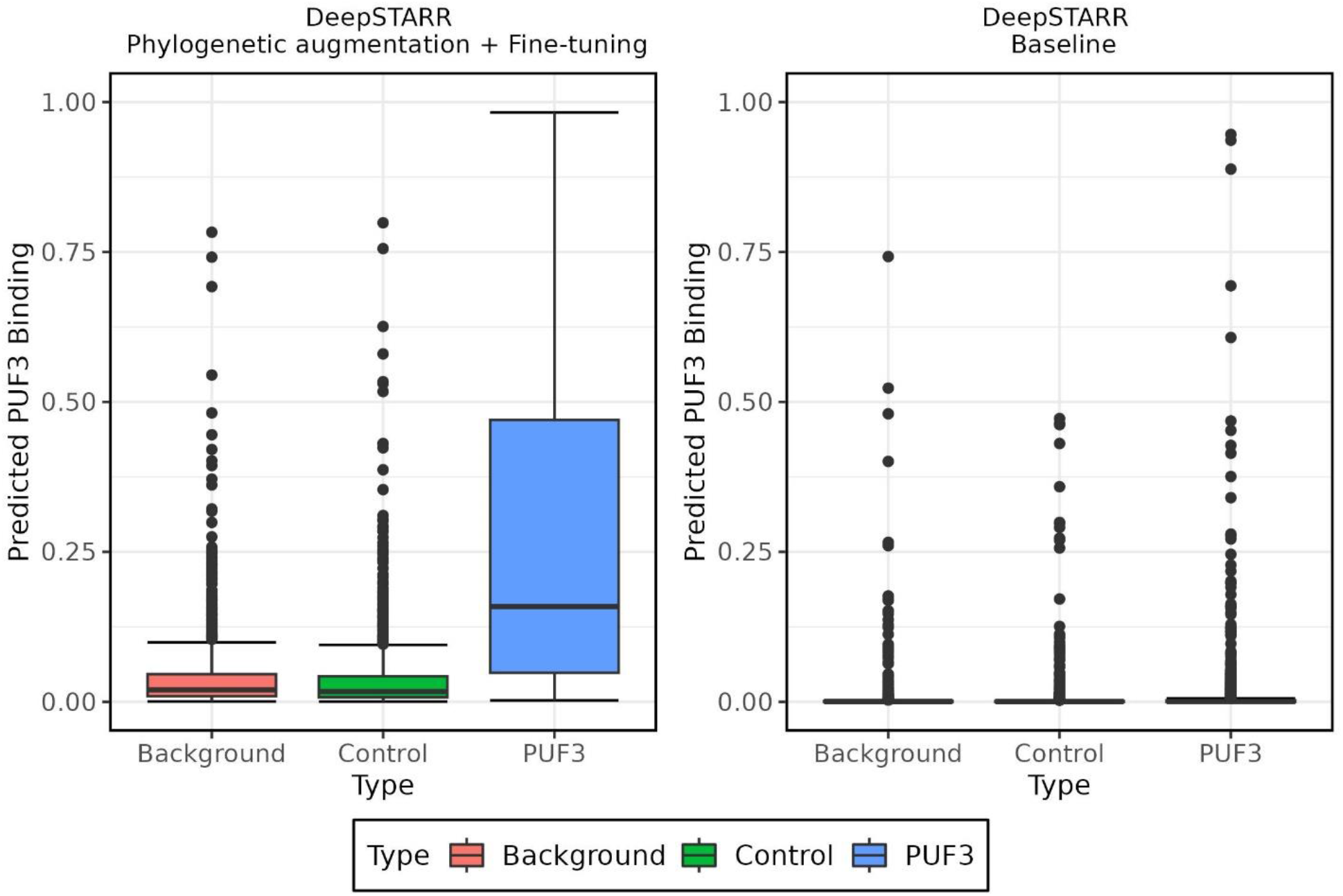
Global importance analysis of the PUF3 consensus motif in the DeepSTARR models. Predicted PUF3 binding class probability for a DeepSTARR model trained with phylogenetic augmentations and fine-tuning (left) and for a baseline DeepSTARR model (right). The x-axis is the type of sequence set and the y-axis is the predicted PUF3 binding (between 0 and 1). The background type is 1,000 random sequences of length 200bp. The PUF3 type contains the 1,000 background sequences with the PUF3 consensus motif randomly inserted. The control type contains the 1,000 background sequences with a scrambled PUF3 consensus motif randomly inserted.

**Supplementary figure 2.**
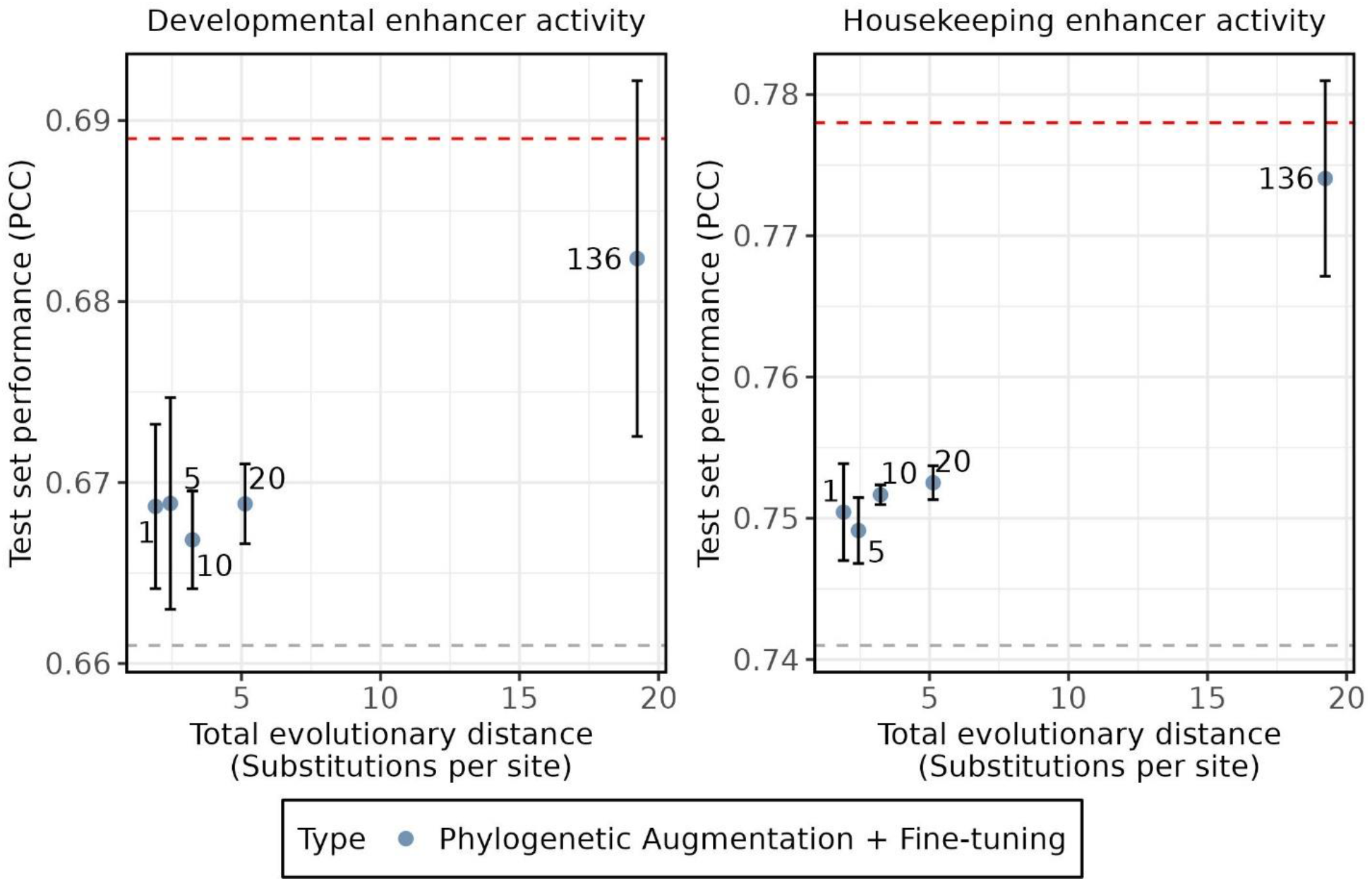
Hyperparameter analysis of species with decreasing evolutionary distance. DeepSTARR test performance (Pearson correlation coefficient; PCC) is shown on the y-axis for *Drosophila* S2 Developmental (left) and Housekeeping (right) enhancer activity for trained models. The x-axis represents the total evolutionary distance of the species used during phylogenetic augmentation with *D. melanogaster*. The labels represent the total number of species used to train each model. The blue dots represent the average test set performance across replicates. The black error bars represent the standard deviation of the three replicates that were trained for each number of species. The dotted grey line represents the average test performance on the original training data with no phylogenetic augmentation or fine-tuning. The dotted red line represents the average test performance on the original training data with phylogenetic augmentation and fine-tuning using all 136 *Drosophila* species and a phylogenetic augmentation rate of 1.

**Supplementary figure 3.**
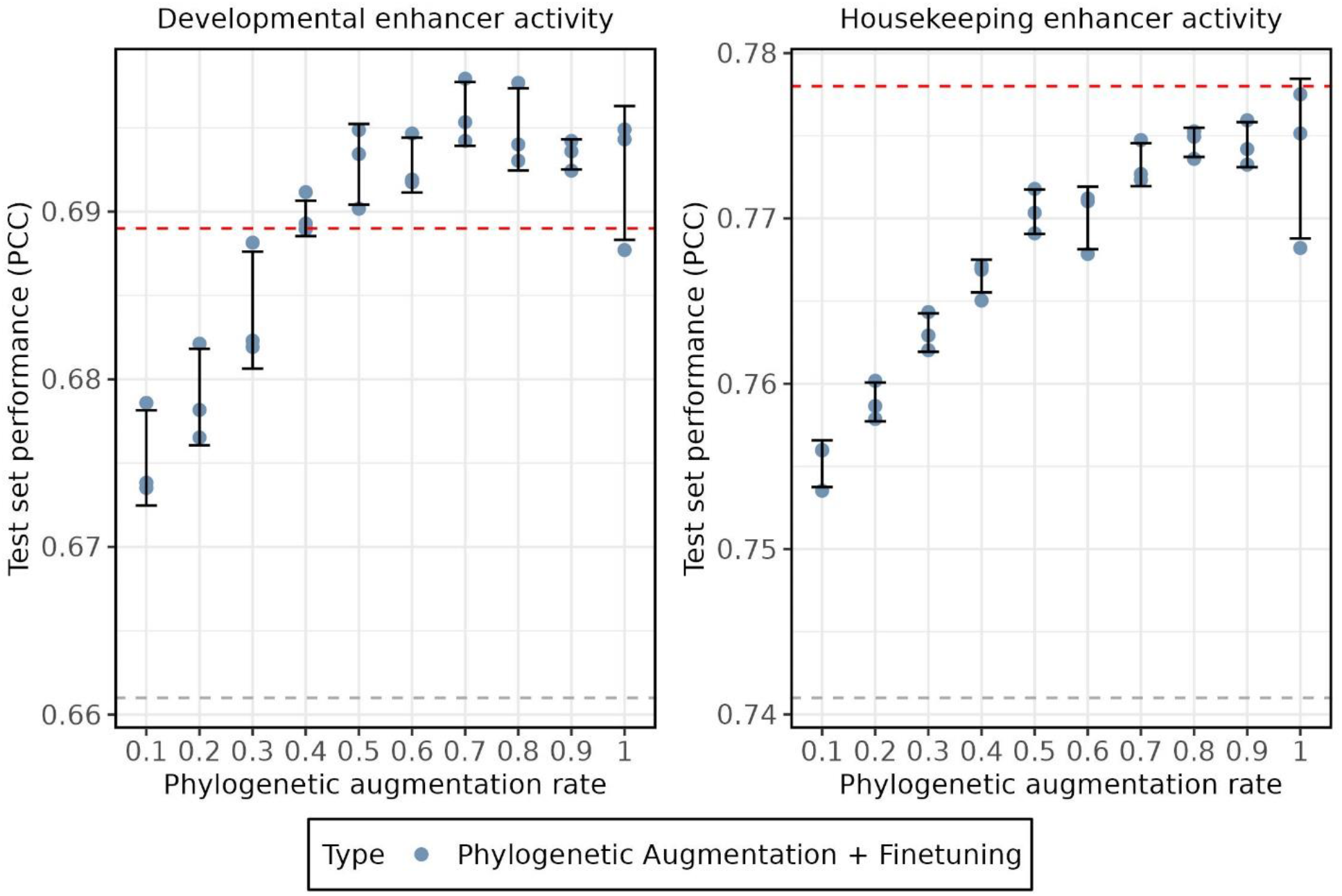
Hyperparameter analysis of phylogenetic augmentation rate with 10 closely related species. DeepSTARR test performance (Pearson correlation coefficient; PCC) is shown on the y-axis for *Drosophila* S2 Developmental (left) and Housekeeping (right) enhancer activity for trained models. The x-axis represents the rate at which phylogenetic augmentation is applied during model training. The blue dots represent the test set performance for individual replicates. The black error bars represent the standard deviation of the three replicates that were trained for each number of species. The dotted grey line represents the average test performance on the original training data with no phylogenetic augmentation or fine-tuning. The dotted red line represents the average test performance on the original training data with phylogenetic augmentation and fine-tuning using all 136 *Drosophila* species and a phylogenetic augmentation rate of 1.

## Acknowledgements

We thank Cameron Dufault, Nawrah Khader, Denise Le, Shanelle Mullany, Ami Sangster, and Tiegh Taylor for comments on the manuscript. We thank Dr. Bernard Kim for sharing the 168-way *Drosophila* genome alignment and Aqsa Alam for sharing the *S. cerevisiae* PUF3 binding data. We thank Dr. Peter Koo for inspiring discussions and comments on the manuscript and an anonymous reviewer for the suggestion to include analysis of untranslated regions.

This work was supported by the Natural Sciences and Engineering Research Council of Canada (NSERC 2020-05972, discovery grant held by J.A.M. and an NSERC discovery grant to A.M.M) This research was performed on infrastructure obtained with grants to A.M.M from the Canada Foundation for Innovation.

## Data availability

The data underlying this article are available on Zenodo. The *Drosophila* multi species genome alignment that we used for this manuscript is available upon request from Dr. Bernard Kim (Stanford University). Additionally, a newer version of the *Drosophila* multi species genome alignment is available that includes additional species^45^ (Supplementary table 5).

## References

1. Kelley, D. R., Snoek, J. & Rinn, J. L. Basset: learning the regulatory code of the accessible genome with deep convolutional neural networks. Genome Res. 26, 990–999 (2016).

2. Avsec, Ž. et al. Effective gene expression prediction from sequence by integrating long-range interactions. Nat. Methods 18, 1196–1203 (2021).

3. Minnoye, L. et al. Cross-species analysis of enhancer logic using deep learning. Genome Res. 30, 1815–1834 (2020).

4. Avsec, Ž. et al. Base-resolution models of transcription-factor binding reveal soft motif syntax. Nat. Genet. 53, 354–366 (2021).

5. de Almeida, B. P., Reiter, F., Pagani, M. & Stark, A. DeepSTARR predicts enhancer activity from DNA sequence and enables the de novo design of synthetic enhancers. Nat. Genet. 54, 613–624 (2022).

6. Koo, P. K. & Ploenzke, M. Deep learning for inferring transcription factor binding sites. Curr. Opin. Syst. Biol. 19, 16–23 (2020).

7. Shrikumar, A. et al. Technical Note on Transcription Factor Motif Discovery from Importance Scores (TF-MoDISco) version 0.5.6.5. (2018) doi:10.48550/ARXIV.1811.00416.

8. Koo, P. K., Majdandzic, A., Ploenzke, M., Anand, P. & Paul, S. B. Global importance analysis: An interpretability method to quantify importance of genomic features in deep neural networks. PLOS Comput. Biol. 17, e1008925 (2021).

9. Novakovsky, G., Fornes, O., Saraswat, M., Mostafavi, S. & Wasserman, W. W. ExplaiNN: interpretable and transparent neural networks for genomics. Genome Biol. 24, 154 (2023).

10. Maslova, A. et al. Deep learning of immune cell differentiation. Proc. Natl. Acad. Sci. 117, 25655–25666 (2020).

11. Abadi, M. et al. TensorFlow: A system for large-scale machine learning. (2016) doi:10.48550/ARXIV.1605.08695.

12. Paszke, A. et al. PyTorch: An Imperative Style, High-Performance Deep Learning Library. (2019) doi:10.48550/ARXIV.1912.01703.

13. De Boer, C. G. et al. Deciphering eukaryotic gene-regulatory logic with 100 million random promoters. Nat. Biotechnol. 38, 56–65 (2020).

14. Tareen, A. & Kinney, J. B. Biophysical models of cis-regulation as interpretable neural networks. (2020) doi:10.48550/ARXIV.2001.03560.

15. de Boer, C. G. & Taipale, J. Hold out the genome: a roadmap to solving the cis-regulatory code. Nature 625, 41–50 (2024).

16. Shorten, C. & Khoshgoftaar, T. M. A survey on Image Data Augmentation for Deep Learning. J. Big Data 6, 60 (2019).

17. Li, B., Hou, Y. & Che, W. Data augmentation approaches in natural language processing: A survey. AI Open 3, 71–90 (2022).

18. Cao, Z. & Zhang, S. Simple tricks of convolutional neural network architectures improve DNA-protein binding prediction. Bioinforma. Oxf. Engl. 35, 1837–1843 (2019).

19. Toneyan, S., Tang, Z. & Koo, P. K. Evaluating deep learning for predicting epigenomic profiles. Nat. Mach. Intell. 4, 1088–1100 (2022).

20. Lee, N. K., Tang, Z., Toneyan, S. & Koo, P. K. EvoAug: improving generalization and interpretability of genomic deep neural networks with evolution-inspired data augmentations. Genome Biol. 24, 105 (2023).

21. Weirauch, M. T. & Hughes, T. R. Conserved expression without conserved regulatory sequence: the more things change, the more they stay the same. Trends Genet. TIG 26, 66–74 (2010).

22. Zoonomia Consortium. A comparative genomics multitool for scientific discovery and conservation. Nature 587, 240–245 (2020).

23. Lu, A. X., Lu, A. X. & Moses, A. Evolution Is All You Need: Phylogenetic Augmentation for Contrastive Learning. (2020) doi:10.48550/ARXIV.2012.13475.

24. Lu, A. X. et al. Discovering molecular features of intrinsically disordered regions by using evolution for contrastive learning. PLOS Comput. Biol. 18, e1010238 (2022).

25. Kelley, D. R. Cross-species regulatory sequence activity prediction. PLOS Comput. Biol. 16, e1008050 (2020).

26. Alam, A., Duncan, A. G., Mitchell, J. A. & Moses, A. M. Functional similarity of non-coding regions is revealed in phylogenetic average motif score representations. http://biorxiv.org/lookup/doi/10.1101/2023.04.09.536185 (2023) xdoi:10.1101/2023.04.09.536185.

27. Hogan, G. J., Brown, P. O. & Herschlag, D. Evolutionary Conservation and Diversification of Puf RNA Binding Proteins and Their mRNA Targets. PLoS Biol. 13, e1002307 (2015).

28. Visel, A., Minovitsky, S., Dubchak, I. & Pennacchio, L. A. VISTA Enhancer Browser--a database of tissue-specific human enhancers. Nucleic Acids Res. 35, D88–92 (2007).

29. Villar, D. et al. Enhancer evolution across 20 mammalian species. Cell 160, 554–566 (2015).

30. She, X. et al. Definition, conservation and epigenetics of housekeeping and tissue-enriched genes. BMC Genomics 10, 269 (2009).

31. Lewin, H. A. et al. Earth BioGenome Project: Sequencing life for the future of life. Proc. Natl. Acad. Sci. 115, 4325–4333 (2018).

32. Sayers, E. W. et al. Database resources of the national center for biotechnology information. Nucleic Acids Res. 50, D20–D26 (2022).

33. Armstrong, J. et al. Progressive Cactus is a multiple-genome aligner for the thousand-genome era. Nature 587, 246–251 (2020).

34. Hickey, G., Paten, B., Earl, D., Zerbino, D. & Haussler, D. HAL: a hierarchical format for storing and analyzing multiple genome alignments. Bioinforma. Oxf. Engl. 29, 1341–1342 (2013).

35. Kuderna, L. F. K. et al. Identification of constrained sequence elements across 239 primate genomes. Nature (2023) doi:10.1038/s41586-023-06798-8.

36. Feng, S. et al. Dense sampling of bird diversity increases power of comparative genomics. Nature 587, 252–257 (2020).

37. Gerber, A. P., Herschlag, D. & Brown, P. O. Extensive association of functionally and cytotopically related mRNAs with Puf family RNA-binding proteins in yeast. PLoS Biol. 2, E79 (2004).

38. Giudice, G., Sánchez-Cabo, F., Torroja, C. & Lara-Pezzi, E. ATtRACT-a database of RNA-binding proteins and associated motifs. Database J. Biol. Databases Curation 2016, baw035 (2016).

39. Zhang, X., Kaplow, I. M., Wirthlin, M., Park, T. Y. & Pfenning, A. R. HALPER facilitates the identification of regulatory element orthologs across species. Bioinforma. Oxf. Engl. 36, 4339–4340 (2020).

40. Zerbino, D. R. et al. Ensembl 2018. Nucleic Acids Res. 46, D754–D761 (2018).

41. Yates, A. et al. The Ensembl REST API: Ensembl Data for Any Language. Bioinforma. Oxf. Engl. 31, 143–145 (2015).

42. Kingma, D. P. & Ba, J. Adam: A Method for Stochastic Optimization. Preprint at http://arxiv.org/abs/1412.6980 (2017).

43. Paten, B. et al. Cactus: Algorithms for genome multiple sequence alignment. Genome Res. 21, 1512–1528 (2011).

44. Huerta-Cepas, J., Serra, F. & Bork, P. ETE 3: Reconstruction, Analysis, and Visualization of Phylogenomic Data. Mol. Biol. Evol. 33, 1635–1638 (2016).

45. Kim, B. Y. et al. Single-fly assemblies fill major phylogenomic gaps across the Drosophilidae Tree of Life. http://biorxiv.org/lookup/doi/10.1101/2023.10.02.560517 (2023) xdoi:10.1101/2023.10.02.560517.

